# A comparative analysis of acellular versus red cell based subnormothermic machine perfusion in human kidney transplantation

**DOI:** 10.64898/2026.01.06.697890

**Authors:** HVM Spiers, ML Nicholson, S Deffrennes, S MacMillan, AL Paterson, M Larraz, I Mohorianu, V Kosmoliaptsis, SA Hosgood

**Affiliations:** Department of Transplantation, Addenbrooke’s Hospital, Cambridge, UK; Department of Surgery, University of Cambridge, Cambridge, UK; NIHR Blood and Transplant Research Unit in Organ Donation and Transplantation at the University of Cambridge, Cambridge, UK; Department of Development and Regeneration, Katholieke Universiteit Leuven, Leuven, Belgium; Department of Nephrology, Dialysis and Renal Transplantation, University Hospitals Leuven, Leuven, Belgium; Department of Histopathology, Addenbrooke’s Hospital, Cambridge, UK; Cambridge Stem Cell Institute, University of Cambridge, UK; Institute of Metabolic Sciences, University of Cambridge, UK

**Keywords:** Normothermic machine perfusion, subnormothermic machine perfusion, acellular perfusate, RNAseq

## Abstract

Normothermic machine perfusion (NMP) typically uses red blood cell (RBC)-based perfusates, with associated challenges, including free haem-driven ferroptotic tubular injury. Acellular perfusates avoid these risks. Subnormothermic machine perfusion (SMP, 32°C) may reduce metabolic demands while preserving organ viability, but the effect of perfusate type under these conditions remains unclear. In a paired experimental design, kidneys from eight deceased donors were randomized 1:1 to 6 hours of SMP with RBC-based (RBC-SMP) or acellular perfusate (SNAP), followed by 4 hours of RBC-based NMP (37°C) to simulate reperfusion. Upregulation of transcriptomic ischaemia-reperfusion injury (IRI) related pathways and downregulation of oxidative phosphorylation at 6 hours SMP and 4 hours reperfusion were similar between RBC-SMP and SNAP kidneys. No differences were observed between groups in transcriptional profiles, metabolic pathway scores, urinary tubular injury biomarker concentrations, or histology. We integrated publicly available NMP transcriptomic data for comparison, demonstrating that compared to 6 hours RBC-NMP at 37°C, both SMP groups showed equivalent pro-inflammatory pathway activation and oxidative phosphorylation depletion. Thus, we provide evidence that SNAP maintains kidneys with equivalent transcriptional and urinary tubular injury biomarker profiles to RBC-SMP. These findings support the feasibility of SNAP for clinical translation in future studies.

## Introduction

Machine perfusion (MP) of deceased-donor kidneys provides advantages compared to static cold storage (SCS) as a method of organ preservation, offering potential for improved clinical outcomes and increased utilisation of organs that may have been previously discarded(1,2). Normothermic machine perfusion (NMP) allows maintenance of organs in a near physiological state and can facilitate organ assessment, repair and regeneration(3–6). The feasibility and safety of NMP in clinical kidney transplantation has been confirmed at the randomised controlled trial level, albeit without yet demonstrating superiority in clinical outcomes compared to traditional SCS (7). Accordingly, unlike in liver transplantation, kidney NMP has not seen widespread clinical implementation, despite renal transplantation representing the most performed solid organ transplant procedure.

NMP typically employs a red blood cell (RBC) based perfusate to deliver oxygen and nutrients during perfusion(3,8). However, RBC based perfusate carries logistical challenges, including requirement for blood type compatibility, infection risk, immunogenicity and haemolysis during perfusion(9,10). Haemolysis increases during kidney NMP, resulting in renal exposure to high levels of free haemoglobin, driving cellular accumulation of haemolysis-derived iron, and subsequent ferroptotic cell death(11), in the absence of the organ’s ability to convert free haemoglobin to bilirubin and excrete it into bile. The problem is exacerbated by the fact that packed RBC units used in kidney NMP are often towards the end of their shelf life and, therefore, carry increased concentrations of free haem (12). These concerns have prompted interest in developing cell-free alternatives, with evidence suggesting that current state-of-the-art extracorporeal membrane oxygenation is efficient enough to consistently achieve physiological and supraphysiological oxygen tension in these solutions, potentially removing the need for an oxygen carrier(13,14). Cell free, or acellular, perfusates could overcome logistical challenges, enhance safety and improve regulatory compliance for widespread implementation of NMP in the clinical setting.

Perfusion at normothermia (37°C) maintains organs in a near physiological state, allowing viability testing(3). A few studies have explored machine perfusion at lower temperatures (e.g. 32°C) in an effort to reduce graft injury whilst maintaining cellular metabolism to a level where oxygen carriers are not required for adequate oxygenation; however, results have been conflicting. Adams et al found that subnormothermic RBC-based machine perfusion (SMP, 32°C) is inferior to NMP(15)(15), raising concerns that lowering perfusion temperature may be detrimental in the context of RBC-based perfusion. Brasile et al demonstrated reduced reperfusion injury and cytoprotective gene induction in a canine DCD kidney transplant model, following perfusion at 32°C with an acellular perfusate, supporting the hypothesis that RBCs may be detrimental in this setting(16). We have recently demonstrated that acellular perfusion at 32°C can maintain kidneys metabolically and increase preservation times, although there was a trend towards mild injury after prolonged preservation (over 12 hours)(13).

Currently, there are no head-to-head comparisons between acellular versus RBC-based kidney machine perfusion at subnormothermic (32°C) temperatures. Investigating the hypothesis that acellular submormothermic perfusion is non inferior to RBC-based perfusion could provide an important milestone in advancing the field. To address this, we employed a same donor, paired kidney (n=8) experimental design, to perfuse kidneys with either RBC or acellular perfusion solution during SMP, followed by a normothermic reperfusion phase using RBC-based perfusion to mimic revascularisation in the recipient. We used untargeted bulk RNAseq transcriptomics to examine gene expression changes between perfusate groups, additionally exploring urinary markers of tubuloepithelial cell injury. We then functionally integrated publicly available bulk RNAseq datasets to explore transcriptomic differences between kidneys undergoing NMP and SMP with RBC and acellular perfusates, to provide insights into pro-inflammatory pathway induction in different machine perfusion contexts.

## Methods

### Study population

Sixteen human donor kidneys from eight deceased donors were recruited into this study after being declined for transplantation and consented for research. Written consent for the use of kidneys for research was given by the donor families and was obtained by Specialist Nurses in Organ Transplantation. Ethical approval was obtained from NRES: 22/WA/0167.

### Study design

Paired kidneys (both kidneys from the same donor) were randomly assigned to undergo acellular or RBC based machine perfusion at 32°C for six hours. Following six hours of SMP, both kidneys were ‘reperfused’ using an RBC-based perfusate at normothermia (37°C), to mimic implantation and reperfusion in a transplant setting (FigS1).

### Machine perfusion

All kidneys were received on ice at 4°C after a period of static cold storage, prepared for machine perfusion and weighed. The renal artery was isolated and kidneys flushed with 500mls of cold Ringer’s solution. The ureter was also cannulated prior to perfusion. All kidneys were perfused for 6 hours at 32°C using an adapted paediatric cardiac bypass system (Medtronic, Biconsole 560), after which they were removed from the circuit and flushed with 1L of cold Ringer’s solution via the renal artery, before a fresh circuit was used to perfuse at 37°C for a further 4 hours. For all perfusions, the perfusion solution was oxygenated (95% oxygen/5% CO_2_) at a flow rate of 0.1L/min, and perfusion was undertaken continuously through the renal artery. Cortical wedge biopsies were taken at selected timepoints with one formalin-fixed (10% formaldehyde) and paraffin-embedded (FFPE) and one stored in RNA*later* for 24 hours prior to transfer to -80°C until analysed. Urine was sampled hourly and the volume replaced with Ringer’s solution; urine samples were centrifuged at 1600RPM for 10min at 4°C, the supernatant (bar 200uL above the buffy coat) was collected and snap frozen in liquid nitrogen then stored at -80°C until analysis.

#### RBC based perfusate

Where RBC perfusate was used (in the RBC arm of the subnormothermic 6 hr perfusion, and for all kidneys in the normothermic reperfusion phase), the circuit was primed with 300ml Ringer’s solution (Baxter Healthcare, Thetford, UK), 15ml Mannitol 10% (Baxter Healthcare), 27 ml sodium bicarbonate 8.4% (Fresenius Kabi, Runcorn, UK), 3000iu heparin (LEO Pharma A/S, Ballerup, Denmark) and 6.6mg Dexamethasone (Hameln Pharmaceuticals, Hamelin, Germany). A unit of ABO compatible packed red cells was then added. Fifteen milliliters of sodium bicarbonate 8.4% (B Braun, Melsungen, Germany) were added. Insulin (100IU; Actrapid, Novo Nordisk, London, UK) was infused at a rate of 20ml/h, in addition to glucose 5% (Baxter Healthcare) at a rate of 5ml/h, and Synthamin 17 10% (Baxter Healthcare, Thetford, UK). Glyceryl trinitrate (10ml of 50mg/ml in 90ml Saline) was also infused at 10ml/hr.

#### Acellular perfusate

Kidneys randomised to the acellular arm during the 6 hours subnormothermic perfusion, were perfused with a Human serum albumin (5%, 250ml) and Ringer’s solution (219ml) based perfusate, supplemented with 6.6mg dexamethasone (Hameln Pharmaceuticals, Hamelin, Germany), 5ml calcium gluconate 10% and 15ml sodium bicarbonate 8.4% (Fresenius Kabi, Runcorn, UK), 250mg meropenem and 2.5mg verapamil. During perfusion Synthamin 17 10% (Baxter Healthcare, Thetford, UK) supplemented with 15mls of sodium bicarbonate 8.4% (B Braun, Melsungen, Germany), 5ml multivitamins, and 1ml insulin (100IU; Actrapid, Novo Nordisk, London, UK) was infused at 10ml/hr. Glucose 5% (Baxter Healthcare), at a rate of 3ml/h, and Glyceryl trinitrate (10ml of 50mg/ml in 90ml Saline) at 10ml/hr for the first hour then 5ml/hr thereafter, were also infused.

### Calculation of oxygen consumption and extraction ratios

#### RBC-NMP

Arterial oxygen content CaO_2_ (ml/dL) = (1.34 (mL O_2_/gram Hb) x Hb (g/dL) x SaO_2_ x 0.01) + (0.023 (mlO_2_/kPa/dL) x PaO_2_ (kPa)) Venous oxygen content CvO_2_ (mL/dL) = (1.34 (mL O_2_/gram Hb) x Hb (g/dL) x SvO_2_ x 0.01) + (0.023 x PvO_2_ (kPa))

#### SNAP

Arterial oxygen content CaO_2_ (mL/dL) = 0.023 (ml O_2_/kPa/dL) x PaO_2_ (kPa) Venous oxygen content CvO_2_ (mL/dL) = 0.023 (ml O_2_/kPa/dL) x PvO_2_ (kPa) Oxygen delivery DO_2_ (ml/min) = (CaO_2_ x 0.01) x flow rate Oxygen consumption VO_2_ (ml O2/min/100g) = (CaO_2_ – CvO_2_) x 0.01 x flow rate Oxygen extraction ratio (OER) = VO_2_/DO_2_

#### Reperfusion 37°C

Arterial oxygen content CaO_2_ (ml/dL) = (1.34 x Hb x SaO_2_ x 0.01) + (0.021 x PaO_2_) Venous oxygen content CvO_2_ (ml/dL) = (1.34 x Hb x SvO_2_ x 0.01) + (0.021 x PvO_2_)

### Haematoxylin and eosin (H&E) staining and histological analysis

Clinical grade H&E staining was completed by the Human Research Tissue Bank at Addenbrooke’s Hospital on 4μm FFPE sections. All H&E-stained sections were imaged using an Olympus IX81 inverted microscope with a 10x objective (Olympus Corporation, Tokyo, Japan). A minimum of five fields of view per section were imaged. A consultant renal transplant pathologist (AP) provided an injury report on representative sections from each timepoint of each experiment, blinded to perfusate allocation and perfusion timepoint.

### Tissue RNA extraction

For 7 kidney pairs, samples were thawed on ice, removed from storage solution and macerated on ice using a sterile scalpel. Macerated tissue was transferred to 700uL QIAzol Lysis Reagent (Qiagen, Hilden, Germany) and homogenised using a sterile pestle followed by 10x passages through a 20g sterile needle. 140uL of chloroform was added and sample shaken for 15 seconds by hand before being centrifuged at 12,000g for 15 minutes at 4°C. The clear upper supernatant (aqueous phase) containing RNA was taken forward and 1.5x the volume of 100% ethanol was added. This was then loaded onto Qiagen RNeasy Midi columns and total RNA extracted according to manufacturer’s instruction. RNA was eluted in 50uL of RNAse free water. DNA was removed on column using the Qiagen RNase-Free DNase set. Quality of RNA was assessed using nanodrop spectrophotometer and an RNA nano Bioanalyzer kit (Agilent Technologies, Palo Alto, CA, USA) using a Bioanalyzer 2100 (Agilent Technologies, Palo Alto, CA, USA). Samples with a RIN >6 were sent for sequencing.

### Bulk RNA sequencing

RNA was shipped on dry ice to Genewiz Azenta (Oxford, UK) for bulk RNA sequencing. Once received, RNA samples were quantified using Qubit 4.0 Fluorometer (Life Technologies, Carlsbad, CA, USA) and RNA integrity was checked with RNA Kit on Agilent 5300 Fragment Analyzer (Agilent Technologies, Palo Alto, CA, USA). rRNA depletion was performed using QIAGEN FastSelect rRNA HMR Kit (Qiagen, Hilden, Germany). RNA sequencing library preparation was performed with NEBNext Ultra II RNA Library Preparation Kit for Illumina following the manufacturer’s recommendations (NEB, Ipswich, MA, USA). Briefly, enriched RNAs were fragmented for 15 minutes at 94 °C. First strand and second strand cDNA were subsequently synthesized. cDNA fragments were end repaired, adenylated at 3’ends, and universal adapters were ligated to cDNA fragments, followed by index addition and library enrichment with limited cycle PCR. Sequencing libraries were validated using the Agilent Tapestation 4200 (Agilent Technologies, Palo Alto, CA, USA), and quantified using Qubit 2.0 Fluorometer (ThermoFisher Scientific, Waltham, MA, USA) as well as by quantitative PCR (KAPA Biosystems, Wilmington, MA, USA).

The sequencing libraries were multiplexed and loaded on the flowcell on the Illumina NovaSeq 6000 instrument according to manufacturer’s instructions. The samples were sequenced using a 2×150 Pair-End (PE) configuration v1.5. Image analysis and base calling were conducted by the NovaSeq Control Software v1.7 on the NovaSeq instrument. Raw sequence output (.bcl files) generated from Illumina NovaSeq was converted into fastq files and de-multiplexed using Illumina bcl2fastq program version 2.20. One mismatch was allowed for barcode matching.

### Transcriptomic analysis

The raw mRNAseq samples were subjected to preprocessing comprising subsampling without replacement(17) to 75M reads to balance variable sequencing depth in samples. Trimming reads to 100 nucleotides (nts) reduced adapter contamination to within accepted levels (<5% across all samples). The proportion of retained biological signal, before and after each processing step, was assessed using fastQC(18), summarised using multiQC.(19) Poor quality samples, from a technical QC perspective, were designated if they contained a significantly higher overall GC%, or low numbers of unique reads indicating degradation, and subsequently excluded (n=5/42 samples, 11.9%). Mapping was performed using STAR (v2.7.4a)(20), with default parameters against vHg38 of the *H sapiens* genome. The quantification of genes, based on the *H Sapiens* gtf annotation was performed using featureCounts(21). Noise correction at expression-matrix level was performed using *noisyR*(22); the normalisation of gene expression was performed using quantile normalisation.(23) Differential expression analysis was performed using *DESeq2*(24). A log_2_(FC) threshold cut off of 0.75 and adjusted p-value 0.05, using Benjamini-Hochberg multiple testing correction, were used to predict DE mRNA transcripts for subsequent analyses. For gene set enrichment analysis (GSEA), all differentially expressed genes were ranked by the inverse of the p value with the sign of the log fold-change, then ran against the Hallmark’s database within MSigDB(25), using the GSEA tool from the Broad Institute(26) with the pre-ranked option, and the following parameters: *No_collapse, classic, default settings*. Single sample GSEA (*ssGSEA*) was carried out using normalized counts from quantile normalization using the GenePattern platform(27).

### Transcriptomic Pathway Activity Scoring

The gene sets used for scoring of glycolysis, fatty acid oxidation, gluconeogenesis, polyol metabolism, tricarboxylic acid cycle, glutamine synthesis, glutathione metabolism and osmotic stress, were obtained from Harmonizome(28). For each sample the average expression of genes from the selected pathway gene set was calculated, and then subtracted with the average expression of another reference set of genes, in which the reference set was randomly sampled from all genes, and the subtracted value used as the “score”(29).

### De Haan et al publicly available transcriptomic data

Raw data were downloaded from GEO GSE289688(14), annotated with relevant metadata provided by the authors, and re-normalised as outlined above.

### ELISA

Assays were performed as directed by the manufacturer instructions. Urinary levels of kidney damage markers neutrophil gelatinase-associated lipocalin (NGAL) and liver fatty acid-binding protein (L-FABP) were measured using the Human Lipocalin-2/NGAL DuoSet ELISA kit (DY1757, R&D Systems) and the Human FABP1/L-FABP DuoSet ELISA kit (DY9465-05, R&D Systems). Urinary levels of proximal tubular endothelial cell damage marker insulin-like growth factor-binding protein 7 (IGFBP7) and distal tubular endothelial cell damage marker tissue inhibitor of metalloproteinases 2 (TIMP2) were measured using the Human IGFBP-rp1/IGFBP-7 DuoSet ELISA set (DY1334-05, R&D Systems) and Human TIMP-2 DuoSet ELISA (DY971, R&D Systems), respectively as per the manufacturers instructions. Absorbance was read at 450nm with 560nm correction on FLUOstar Omega Microplate Reader (BMG LABTECH GmbH) with FLUOstar OPTIMA software (BMG LABTECH GmbH).

### Statistical analysis

Continuous data were tested for normality with the Shapiro–Wilk test. Average values are presented as mean ± standard deviation (SD) for entries following a normal distribution and median with interquartile range (IQR) otherwise. Comparisons between three or more groups were performed using a mixed effects model or Kruskal–Wallis tests with Tukey’s or Dunn’s multiple comparisons corrections respectively. A two-tailed p-value of <0.05 is considered statistically significant. Analyses were conducted in GraphPad PRISM v10.1.1 (San Diego, CA, USA).

## Results

### Haemodynamic, oxygenation and urinary parameters during subnormothermic machine perfusion

We perfused 8 pairs of human kidneys procured for transplantation but subsequently declined and offered for research. Donors had a median age of 65 years (range 45-75), a median BMI of 24.77kg/m^2^ (range 22.76-32.87), were predominantly male (57%) and mostly from brain stem death donation donors (DBD, 71%). The median cold ischaemia time was 28.2 hours (range 13.4-29.6). Full demographic data are presented in Supplementary Table 1.

Kidneys were randomised 1:1 to undergo six hours of either subnormothermic machine perfusion with RBC-based perfusate (RBC-SMP) or subnormothermic machine perfusion with acellular perfusate (SNAP). Following SMP, all kidneys underwent ‘reperfusion’ using RBC based NMP at 37°C for four hours (FigS1). Renal blood flow (RBF) in SNAP kidneys increased significantly during perfusion (Fig1A), rising from a mean of 120.2 ml/min/100g at the start of perfusion, to mean of 194.9 ml/min/100g by 6 hours (p=0.0037). In kidneys perfused with RBC-SMP there was no significant change in RBF across the duration of perfusion (91.11ml/min/100g to 131.8ml/min/100g, p=0.6566). There were no significant differences in RBF between RBC-SMP and SNAP cohorts at any timepoint (Fig1A). Oxygen consumption was similar in both cohorts throughout perfusion (Fig1B), however, the oxygen extraction ratio was significantly higher in SNAP kidneys after 1 hour of perfusion (Fig1C). Taken together, these data suggest SNAP provides adequate tissue oxygen delivery through increased extraction of oxygen compared to RBC-SMP. Urine production during SMP was not significantly different between groups at any timepoint (Fig1D), nor was the osmolarity of the urine produced (Fig1E). During the reperfusion phase, both kidney groups underwent RBC-based NMP at 37°C. Renal blood flow, oxygen consumption and extraction, urine output and osmolality were not different between SNAP and RBC-SMP kidneys during the reperfusion phase (Fig1F-J).

**Figure 1.**
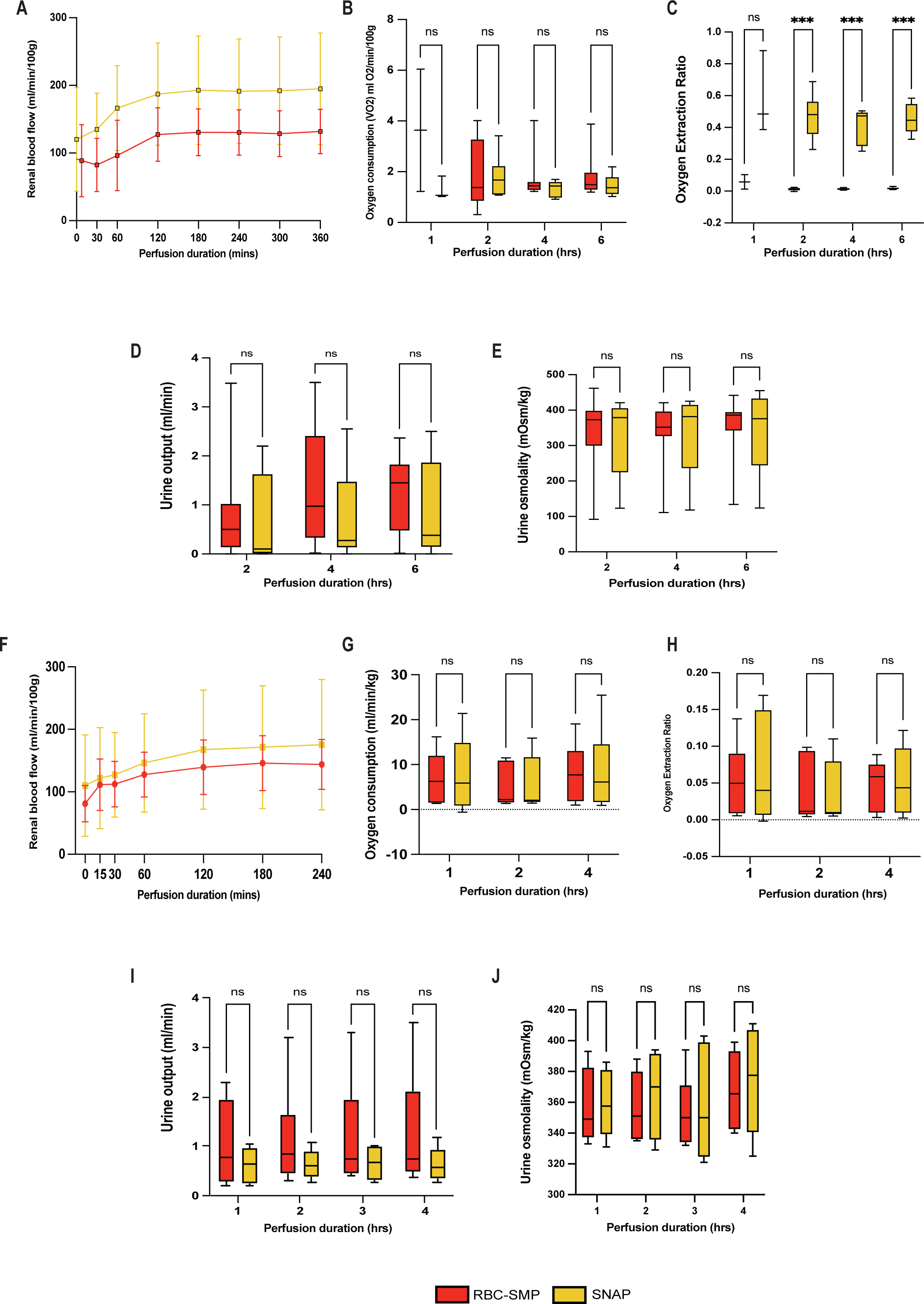
Renal blood flow (**A**), oxygen consumption (**B**) and extraction ratio (**C**), urine output (**D**) and osmolality (**E**) during subnormothermic machine perfusion. Renal blood flow (**F**), oxygen consumption (**G**) and extraction ratio (**H**), urine output (**I**) and osmolality (**J**) during reperfusion with red blood cell based perfusion at 37°C. RBC-SMP, RBC-based subnormothermic machine perfusion; SNAP, subnormothermic acellular perfusion.

### Transcriptional pathways during subnormothermic machine perfusion are similar between RBC-SMP and SNAP kidneys

We investigated tissue transcriptional changes between the two perfusate groups at the end of 6 hours of SMP, and at 4 hours of simulated reperfusion, comparing these to gene expression at the end of SCS (pre-SMP) in our paired kidney experimental design. Principal component analysis (PCA) revealed that the largest source of transcriptional variation was between biopsies taken in the cold (pre-SMP) versus in the warm (SMP and reperfusion timepoints; PC1: 39.76%), highlighting the significant, but anticipated, transcriptional changes upon reperfusion with a warm oxygenated solution following a period of cold preservation (Fig2A). In the top 50 genes accounting for this variation, on PCA loadings, we found transcripts for pro-inflammatory cytokines (*IL1B, IL6, CXCL8*), coagulation pathway modulators (*SERPINE1, PLAU, TFPI2*), activated endothelial cells (*SELE, HBEGF*), oxidative stress and apoptosis regulators (*MT1A,* SLC7A11), and pro-(*MYC, FOSL1*) and anti-inflammatory (*ATF3*) transcription factors (Fig2B). Over-representation analysis of these genes highlighted pathways related to immune activation and inflammatory responses, including cytokine signaling, apoptosis and coagulation (Fig2C), all known to be upregulated in ischaemia-reperfusion injury (IRI)(30–32). PCA also highlighted variation in gene expression between sex groups (PC2: 20.04%) at all timepoints (Fig2D), prompting us to account for this as a covariate in our differential expression analysis. Differential gene expression at six hours of SMP showed that only one gene was significantly different between SNAP and RBC-SMP groups, namely the beta haemoglobin gene *HBB*, which was downregulated in SNAP kidneys (Fig2E). These results highlight transcriptional similarity at tissue level between the two groups, regardless of the perfusate method used during machine perfusion.

**Figure 2.**
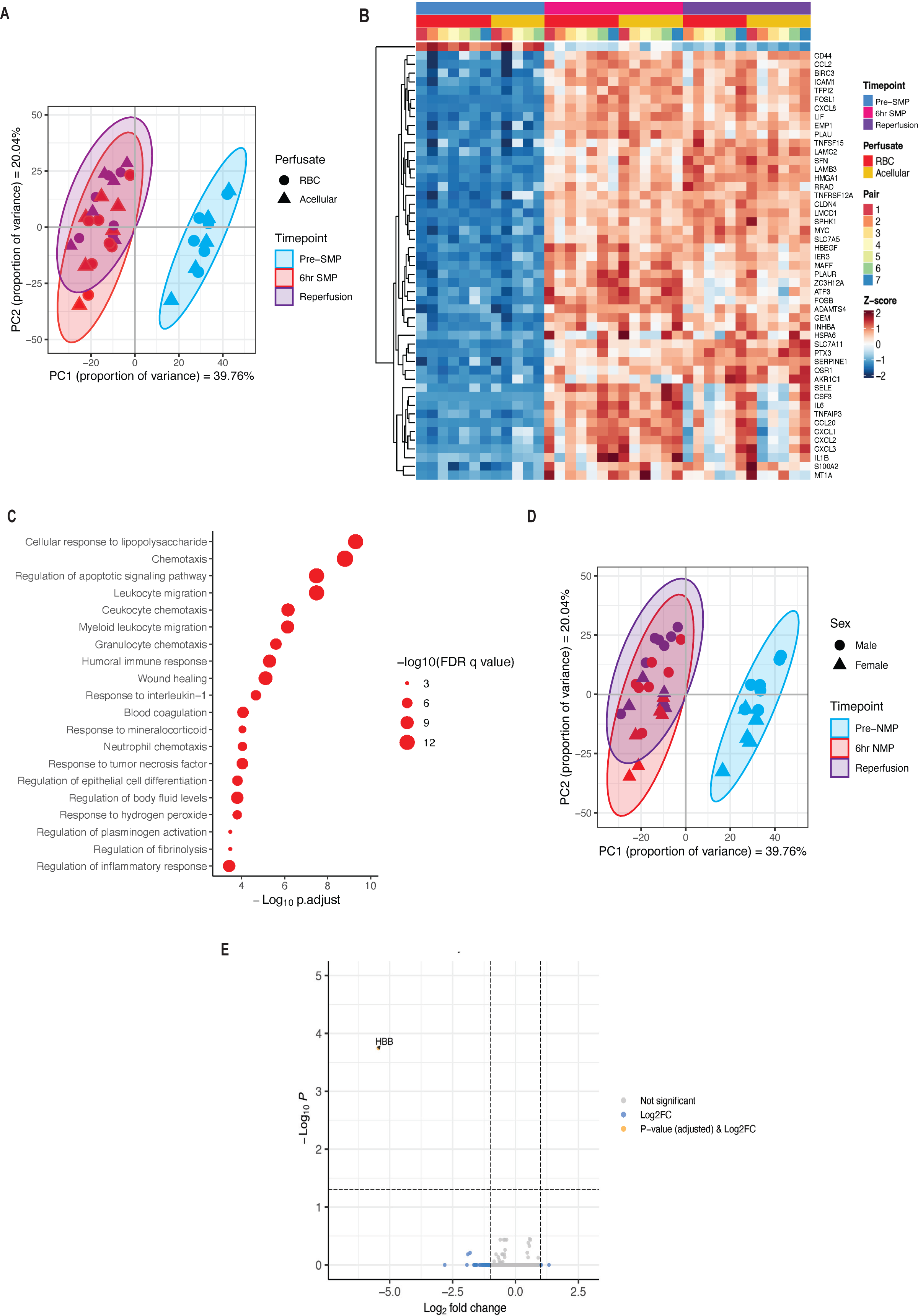
**A**: Assessment of gene expression by principal component analysis (PCA) using the top 500 most variable genes. **B**: Heatmap depiction of the top 50 most variable genes from the PC1 loading. **C**: Reactome pathways annotated from genes in B. **D**: PCA from A annotated for donor sex. **E**: Volcano plot demonstrating only one significant differentially expressed gene between RBC-SMP and SNAP kidneys after 6 hours SMP. RBC-SMP, RBC-based subnormothermic machine perfusion; SNAP, subnormothermic acellular perfusion.

### Changes in pro-inflammatory pathway activation in kidneys during SMP are independent of perfusate used

We next sought to explore the potential impact of the different perfusion methods on global transcriptomic pathway changes during SMP and at 4 hours of simulated reperfusion. Gene set enrichment analysis (GSEA) against the MSigDB Hallmarks pathways(25) revealed enrichment of pro-inflammatory/immune activation and repair/proliferation pathways after 6 hours of SMP (Fig3A). Enriched pathways included *TNFa signaling via NFkB*, *Allograft rejection*, *Inflammatory response* and *Complement*, all of which are known to be central in kidney IRI during machine perfusion at normothermia(30,31). Gene sets relating to cytokine and chemokine signaling were also enriched, including *IL6/JAK/STAT3, IL2 STAT5 Signaling* and *Interferon Gamma Response.* Additionally, there was upregulation of genes annotated to pathways indicative of cell proliferation (*MYC targets)* and *epithelial-mesenchymal-transition*. The enrichment of these pathways was observed to be of similar magnitude between RBC-SMP and SNAP cohorts, both at 6 hours of SMP and at 4 hours of reperfusion. Furthermore, in both perfusate groups there was downregulation of the *Oxidative Phosphorylation* gene set at the end of SMP and at reperfusion, compared to end of SCS.

**Figure 3.**
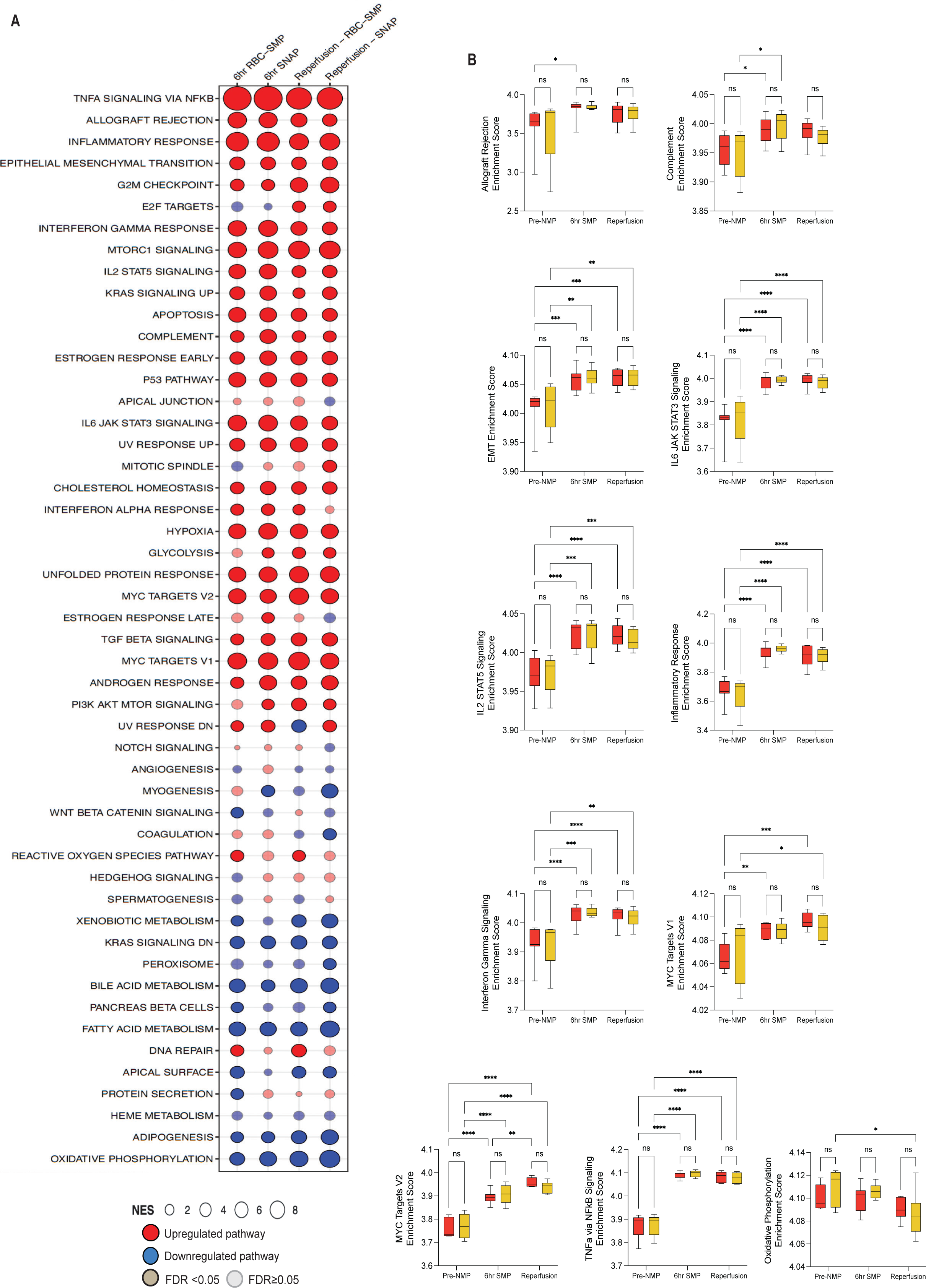
**A**: GSEA against the Hallmarks pathways using differentially expressed genes between pre-SMP timepoint biopsy and annotated timepoint. **B**: Log2 transformed ssGSEA scores and comparisons for select pathways; red represents RBC-SMP and yellow SNAP. RBC-SMP, RBC-based subnormothermic machine perfusion; SNAP, subnormothermic acellular perfusion; FDR, false discovery rate.

Having demonstrated similar patterns of global gene enrichment between SNAP and RBC-SMP cohorts, we used single sample GSEA (ssGSEA) to quantify differences between perfusion groups, both at the end of SMP and at 4 hours of simulated reperfusion. This analysis showed an increase in the enrichment scores for the aforementioned pro-inflammatory/immune activation pathways between the end of SCS to the six hour SMP timepoint, however, there were no significant differences in the magnitude of that enrichment between RBC-SMP and SNAP kidneys (Fig3B). The same was true for group comparisons at the reperfusion phase, with no significant differences between perfusion cohorts. This suggests the perfusate method had no differential effect on the magnitude of the primary IRI affecting all kidneys during SMP, or on the magnitude of the secondary IRI during the simulated reperfusion phase. Interestingly, although ssGSEA confirmed no significant differences, at any timepoint, between RBC-SMP and SNAP kidneys, within group comparisons for the *Oxidative phosphorylation* pathway, demonstrated greater pathway downregulation within SNAP kidneys between SCS and reperfusion timepoints (Fig3B).

We next sought to explore how SMP compares to NMP at the global transcriptional level. We capitalised on publicly available bulk transcriptomic data from the study by de Haan et al(11) to explore transcriptional signatures induced after RBC-NMP at 37°C. GSEA of differentially expressed genes at 6 hours of RBC-NMP, RBC-SMP and SNAP (compared to biopsies at SCS) revealed induction of similar pro-inflammatory and immune activation pathways associated with kidney IRI(30,31), including *TNFalpha via NFkB signaling*, *Inflammatory response*, *Allograft rejection* and *IL6 JAK STAT3 Signalling* (FigS2). This enrichment was of similar magnitude in all groups, suggesting that lowering temperature to 32°C does not mitigate IRI, regardless of RBC-based or acellular perfusate. Notably, depletion of the *Oxidative Phosphorylation* pathway was seen at the 6 hour timepoint in all machine perfusion groups. Taken together, these data suggest that kidneys experience a similar magnitude of IRI at 6 hour NMP and SMP.

### Acellular versus RBC-based perfusate does not impact metabolism-associated transcriptional pathways

A cell-free nutrient-supplemented perfusate has been demonstrated to facilitate extended *ex vivo* preservation of metabolically active kidneys(14). Transcriptomic pathway analysis has been previously used to gain metabolic insights into kidney physiology(33). Based on this precedent, we leveraged our transcriptomic approach to explore changes in several relevant metabolic pathways in kidneys undergoing RBC-SMP and SNAP (see Transcriptomic Pathway Activity Scoring; Methods). We found that for all pathways explored, there was no significant difference between RBC-SMP and SNAP kidneys at any timepoint, suggesting that an acellular perfusate can support kidney metabolic functions during six hours of SMP (Fig4). RBC-SMP kidneys demonstrated an increase in glycolysis scores during SMP and at reperfusion, compared to pre-SMP, whereas SNAP kidneys showed no difference (Fig4A). The tricarboxylic acid (TCA) score gradually reduced during SMP and at reperfusion for RBC-SMP kidneys but was only reduced at reperfusion (compared to the pre-SMP timepoint) in the SNAP cohort (Fig4B). In addition to glucose, glutamine is an important nutrient source for glycolysis and the TCA cycle. In both groups, glutamine synthesis scores significantly increased during SMP, with a trend towards further increase at reperfusion, potentially reflecting utilisation and depletion of this cellular energy source in the period of ischaemia inherent to organ procurement and transport on SCS, followed by repletion during SMP and during reperfusion (Fig4C). Fatty acid oxidation (FAO) pathway scores were seen to be reduced in both groups at four hours of reperfusion (Fig4D), with additional reduction in gluconeogenesis scores (Fig4E); coupled with enhanced glycolysis pathway scores throughout perfusions, this may reflect metabolic reprogramming to compensate for energy shortage during acute kidney injury. Polyol metabolism converts glucose to fructose via sorbitol, typically in hyperglycaemia but also under normoglycaemia in ischaemia/hypoxia, inducing osmotic and oxidative stress; it has been implicated in tubular cell injury during IRI(34,35). In both cohorts, polyol metabolism scores reduced during SMP with a further significant reduction at reperfusion (Fig4F), potentially reflecting restoration of canonical glucose metabolism during oxygenated perfusion.

**Figure 4.**
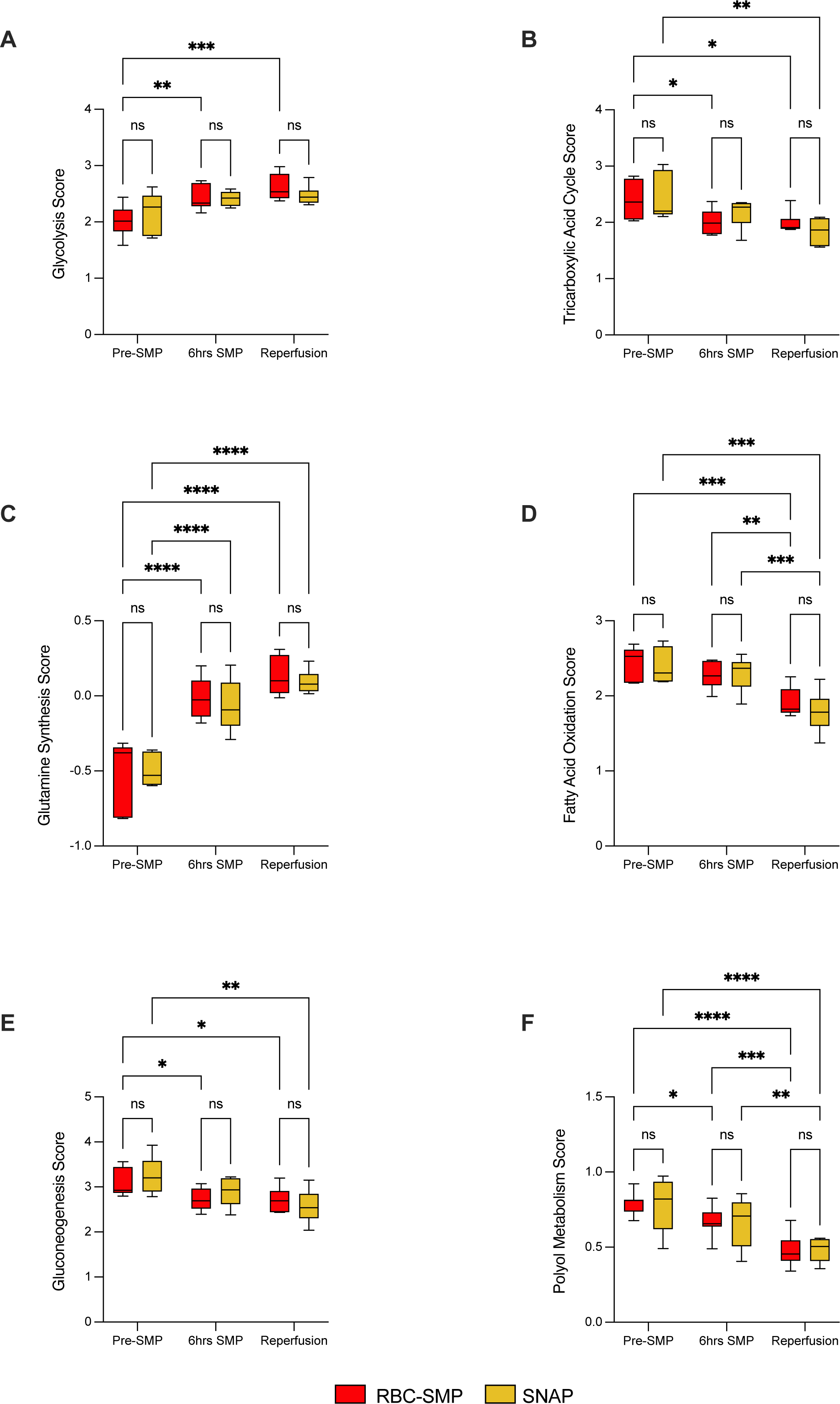
Metabolic pathway scores for glycolysis (**A**), tricarboxylic acid cycle (**B**), glutamine synthesis (**C**), fatty acid oxidation (**D**), gluconeogenesis (**E**) and polyol metabolism (**F**). RBC-SMP, RBC-based subnormothermic machine perfusion; SNAP, subnormothermic acellular perfusion; NES, normalized enrichment score; FDR, false discovery rate.

### Urinary biomarkers of tubuloepithelial cell injury are similar in both RBC-SMP and SNAP perfused kidneys

In homeostasis, there is a significant gradient in oxygen pressure and osmolarity over the corticopapillary axis of the kidney, leading to increased osmotic cellular stress(36). We hypothesised that restoration of metabolic function during SMP may restore osmotic cellular stress. Therefore, we defined an osmotic stress score by averaging expression of related genes (28,33) and assessed changes at 6 hours of SMP and at 4 hours of simulated reperfusion (Fig5A). The osmotic stress score increased in both RBC-SMP and SNAP kidneys during SMP, without further increase at reperfusion, with no significant difference between groups, suggesting an acellular perfusate does not increase osmotic stress relative to RBC based perfusion.

**Figure 5.**
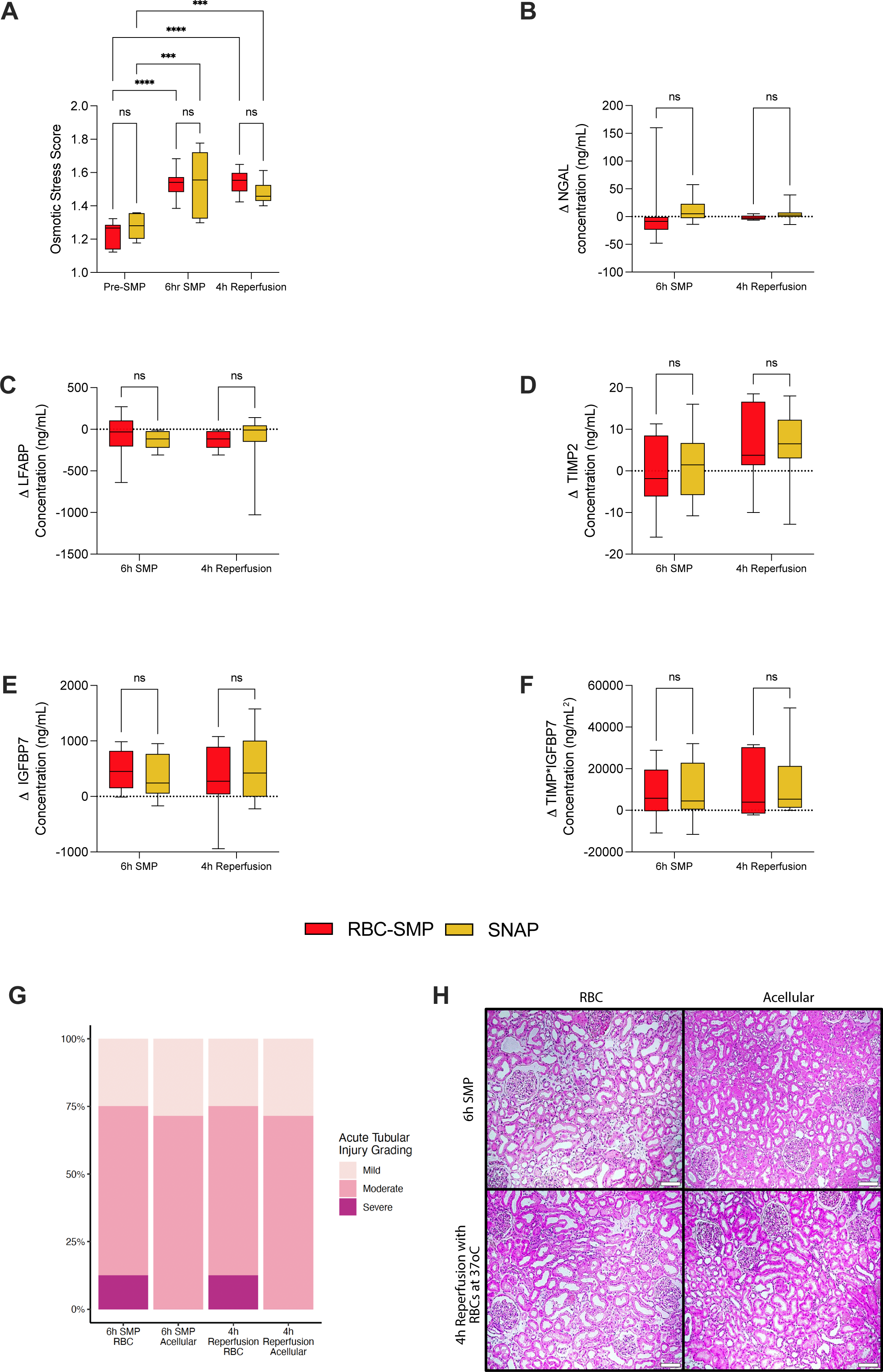
**A**: Osmotic stress score across perfusion timepoints. Change in urine concentration of key urinary injury biomarkers, including NGAL (**B**), L-FABP (**C**), TIMP2 (**D**), IGFBP7 (**E**) and multiplicated TIMP2*IGFBP7 (**F**). **G**: Histological assessment of acute tubular injury grading between perfusion groups. **H**: Representative H&E images of kidneys at 6 hours of SMP and 4 hours of reperfusion. RBC-SMP, RBC-based subnormothermic machine perfusion; SNAP, subnormothermic acellular perfusion.

Having identified an upregulation of osmotic stress associated genes during SMP in both groups, we hypothesised that in the context of pro-inflammatory pathway upregulation (discussed earlier) this may reflect tubuloepithelial cell injury rather than purely homeostatic osmotic stress. Therefore, we sought to explore at the protein level whether there were differences in urinary biomarkers of tubular injury between RBC-SMP and SNAP kidneys. We selected Neutrophil Gelatinase-Associated Lipocalin (NGAL)(37–39), Liver-type Fatty Acid Binding Protein (L-FABP)(40,41), Tissue Inhibitor of Metalloproteinase 2 (TIMP2) and Insulin-like growth factor-binding protein 7 (IGFBP7)(42,43) because of their association with post-transplant outcomes or acute kidney injury during machine perfusion. Additionally, we examined concentrations of combined TIMP and IGFBP7 markers (TIMP*IGFBP7) given their value in predicting adverse outcomes in acute kidney injury(44,45). We assessed the change in urinary concentration of each marker between timepoints (2 hours versus 6 hours SMP; 1 hour versus 4 hours reperfusion). There were no significant differences in the change in urinary concentration of the above urinary injury markers between RBC-SMP and SNAP kidneys during SMP, or during the reperfusion phase. These findings were corroborated by histological assessment by a blinded transplant histopathologist which showed similar tubular injury profiles between RBC-SMP and SNAP kidneys, both during SMP and at reperfusion (Fig5G&H).

## Discussion

Kidney machine perfusion offers advantages over static cold storage (SCS), but commonly used RBC-based perfusates carry logistical challenges and can drive ferroptotic tubular injury via free haem release by RBC lysis. Development of cell free/acellular perfusates and subnormothermic perfusion (32°C) may reduce metabolic demands and mitigate reperfusion injury. In this study, we used a paired kidney experimental design to provide the first direct comparison of RBC and acellular perfusates during subnormothermic machine perfusion of human kidneys at 32°C. We performed bulk RNA sequencing, demonstrating the dominant transcriptomic pathway changes during SMP and at simulated reperfusion with RBCs at 37°C were related to ischaemia–reperfusion injury, including upregulation of *TNFa via NFkB Signaling, Allograft Rejection* and *Inflammatory Response*, and were not different between RBC-SMP and SNAP kidneys. Similarly, the downregulation of *Oxidative phosphorylation* relative to SCS was comparable, with no significant modulation by perfusate type. We explored transcriptional changes in metabolic programs during SMP and at simulated reperfusion, suggesting that SNAP provides equivalent support to select renal metabolic functions as RBC-SMP. We further demonstrated by histological comparison and direct measurement of validated urinary biomarkers of acute tubular injury, that SNAP did not induce more tubuloepithelial injury compared to RBC-SMP. Finally, our analysis integrating publicly available transcriptomic data suggested that RBC-based normothermic machine perfusion induces similar transcriptomic changes compared to subnormothermic perfusion.

A key observation was the similarity in transcriptional changes associated with metabolic pathways between SNAP and RBC-SMP kidneys, with no significant differences between groups at 6 hours of SMP and at simulated reperfusion. This suggests similar patterns of changes in metabolism after cold storage when using an acellular perfusate compared to RBC-based perfusion. Transcription of glutamine synthesis associated genes increased during SMP and at simulated reperfusion, suggesting replacement of this key anaplerotic substrate after ischaemic depletion; prior evidence implicates glutamine in renoprotection during ischaemia-reperfusion injury due to its role in intracellular redox homeostasis(46). Reductions in fatty acid oxidation and gluconeogenesis pathway genes at reperfusion suggested a shift toward glucose utilisation as its availability was restored, while suppression of polyol metabolism implied a move away from stress- and IRI-associated glucose fluxes linked to tubular injury. Interestingly, while RBC-SMP kidneys showed a greater relative induction of glycolysis across the duration of SMP and simulated reperfusion, and earlier reduction of inferred TCA cycle activity, these kinetic differences did not alter overall transcriptional metabolic trajectories, suggesting that oxygen carrier presence is not required for the major transcriptomic features of metabolic recovery at 32°C. The TCA cycle has been previously shown to be maintained during kidney perfusion by an acellular perfusate up to four days of MP (14); the lagging changes in TCA pathway scores seen in SNAP versus RBC-SMP kidneys within our study may reflect differential metabolic maintenance of kidneys *ex vivo* using an acellular perfusate. Future studies combining transcriptomics with metabolomic profiling or isotope tracing are required to validate our findings at tissue level.

We acknowledge the limitations of our study, including modest sample size, reflecting the limited availability of paired human kidneys declined for transplantation. To mitigate this, we employed a paired design, using kidneys from the same donor for both arms, which controls for inter-donor variability and strengthens internal validity. We accept our study did not have the power to assess the impact of donor heterogeneity, including donation type or cold ischaemic time, on the results of our study. Our transcriptomic findings would have been strengthened by validation at the protein level, and this should be included in future work. Nevertheless, we complemented the transcriptomic analysis with validated urinary biomarkers of tubular injury, as well as histological assessment, which provided orthogonal support for comparable IRI in RBC-SMP and SNAP kidneys. Further studies are required to understand the impact of different perfusates at a more granular level and longer perfusion timepoints, including spatial localisation of transcriptomic pathway activation to understand cell specific responses. Finally, these kidneys were not transplanted, so we cannot directly assess *in vivo* graft function. We included a normothermic reperfusion phase to simulate implantation and did not find any differences between acellular and RBC-based perfusion methods.

Taken together, our data support the non-inferiority of SNAP compared to RBC-SMP. Acellular perfusates offer logistical and potential safety advantages, by avoiding the requirement for blood type matching and the risks of haemolysis and free haem toxicity. Our findings provide the foundation for future studies assessing the safety and efficacy of SNAP in the clinical setting.

## Data Availability Statement

The datasets presented in this study can be found in online GEO accession repositories: https://www.ncbi.nlm.nih.gov/geo/

## Acknowledgements

The author(s) declare financial support was received for the research and/or publication of this article. We acknowledge funding support from National Institute for Health and Care Research (NIHR) Blood and Transplant Research Unit in Organ Donation and Transplantation (NIHR203332), a partnership between NHS Blood and Transplant, University of Cambridge and Newcastle University. HS acknowledges funding from a Sir Roy & Lady Calne Royal College of Surgeons Research Fellowship (G121988) and a Wellcome Trust Clinical Research Training Fellowship (G115288). IM acknowledges funding from Wellcome Trust (203151/Z/16/Z) and UKRI Medical Research Council (MC_PC_17230). VK acknowledges funding from an NIHR Fellowship (PDF-2016-09-065) and as a Paul I. Terasaki Scholar (G106170). We would like to acknowledge Maja Kaczmarek and Léonie Walker-Panse for help with perfusion experiments.

## Figure Legends

**Supplementary Figure 1.**

**A**: Schematic of experimental design demonstrating subnormothermic machine perfusion phase and 4 hours reperfusion with RBC-based perfusate at 37°C. Figure produced using Biorender.

**Supplementary Figure 2.**

**A**: GSEA against the Hallmarks pathways using differentially expressed genes between pre-perfusion timepoint biopsy and 6 hour timepoint biopsy for the three perfusion types. Raw GSEA output data available in Supplementary Table 2. RBC-SMP, RBC-based subnormothermic machine perfusion; SNAP, subnormothermic acellular perfusion; RBC-NMP, RBC-based normothermic machine perfusion; NES, normalized enrichment score; FDR, false discovery rate.

