## Supplementary Figure 1 for "A comparative analysis of acellular versus red cell based subnormothermic machine perfusion in human kidney transplantation"

### Slide 1
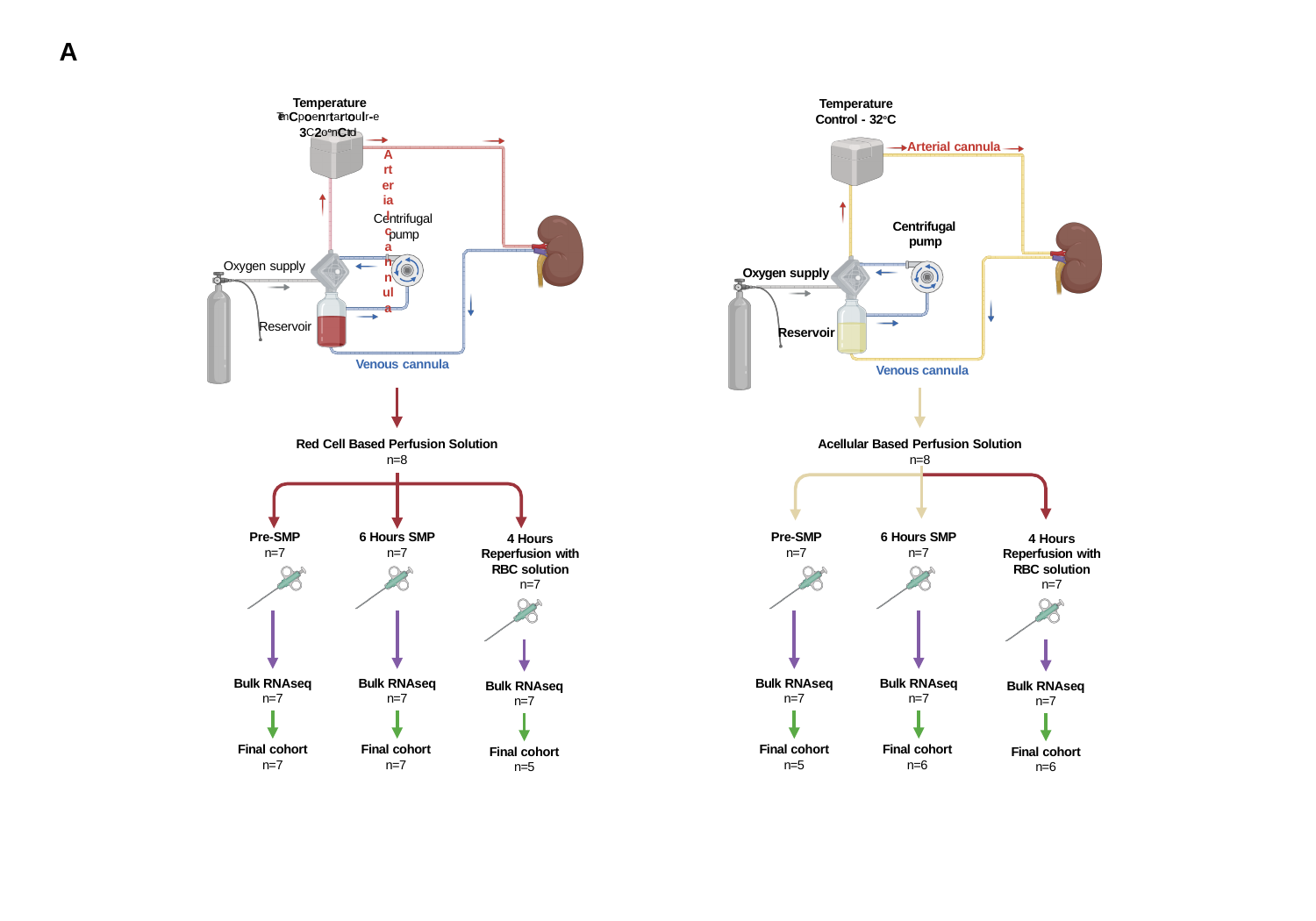

A
Temperature
^Tem^C^p^o^e^n^r^t^a^r^t^o^u^l^r^-^e^ 3^C^2^oon^C^trol^
Arterial cannula
Temperature Control - 32^o^C
Arterial cannula
Centrifugal pump
Centrifugal pump
Oxygen supply
Oxygen supply
Reservoir
Reservoir
Venous cannula
Venous cannula
Acellular Based Perfusion Solution
n=8
Red Cell Based Perfusion Solution
n=8
6 Hours SMP
n=7
Pre-SMP
n=7
6 Hours SMP
n=7
Pre-SMP
n=7
4 Hours Reperfusion with RBC solution n=7
4 Hours Reperfusion with RBC solution n=7
Bulk RNAseq
n=7
Bulk RNAseq
n=7
Bulk RNAseq
n=7
Bulk RNAseq
n=7
Bulk RNAseq
n=7
Bulk RNAseq
n=7
Final cohort
n=7
Final cohort
n=7
Final cohort
n=5
Final cohort
n=6
Final cohort
n=5
Final cohort
n=6
