## Supplementary figures and images for "A comparative analysis of acellular versus red cell based subnormothermic machine perfusion in human kidney transplantation"

### Supplementary Figure 2

#### Slide 1
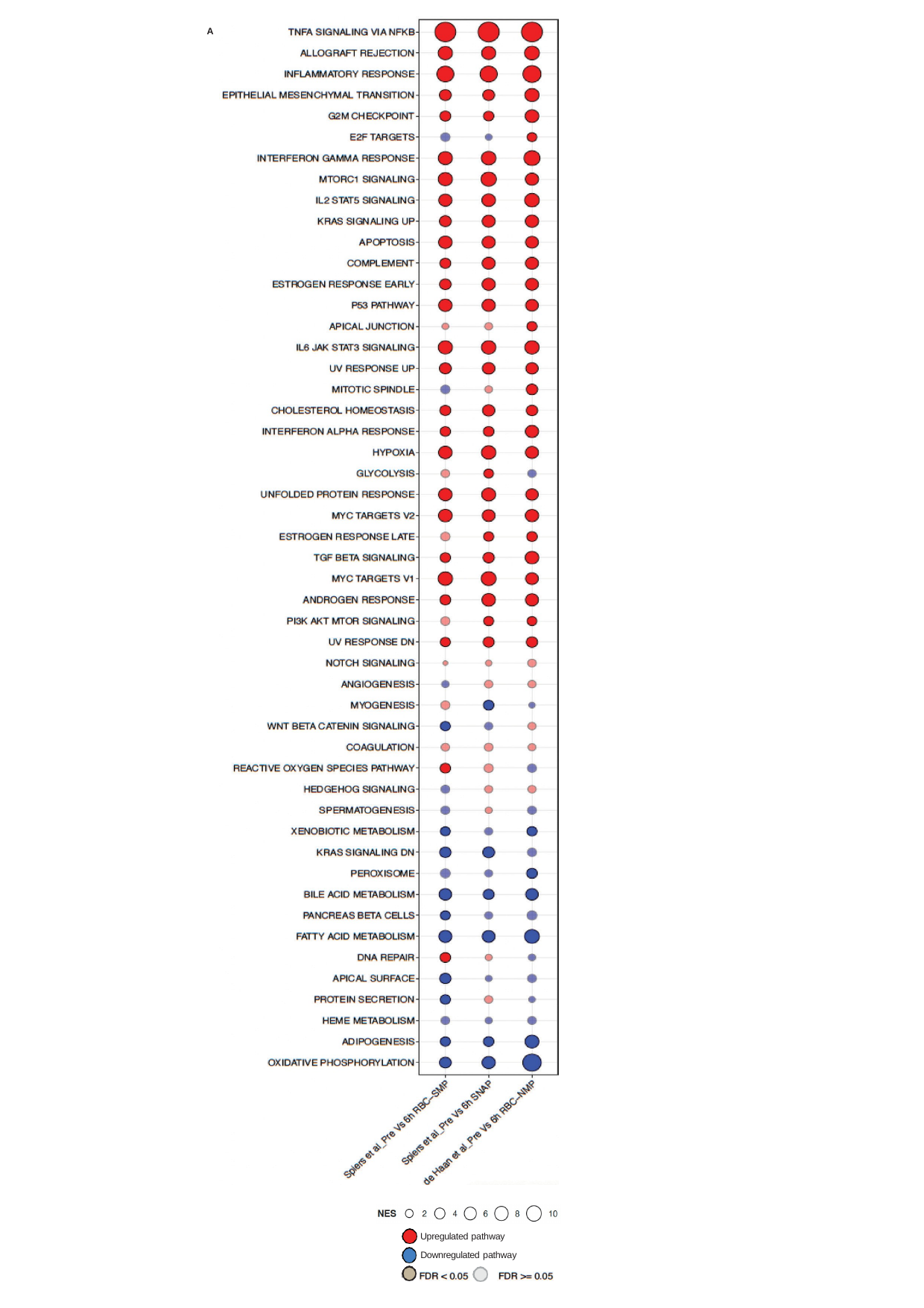

A
Upregulated pathway Downregulated pathway
